# XomicsDB: a database of paired multi-omics datasets for systems biology

**DOI:** 10.1101/2025.09.27.679001

**Authors:** Tianyue Zhang, Li Liu, Yuxuan Wu, Songwook Jung, Xiaoxi Liu, Qizhou Jiang, Ningyuan Yang, Semin Kim, Yi Tao, Gaoyang Li, Huansheng Cao

## Abstract

Life science research is primarily driven by data integration on the basis of theoretical development. Broad applications of multiomics (xomics) data calls for efficient integration of xomics data to infer cellular operation and regulation. To that end, it is urgently needed good-quality xomics data should be curated and shared for the community. Here we present XomicsDB, a database curating paired xomics datasets at both bulk and single-cell levels in model organisms. It contains 183 datasets from 13 model organisms, with a total of seven xomics data types. Users can download these datasets individually or in a batch from the ftp server provided. Users can also upload their paired xomics datasets for sharing. Incoporation of xomics data intom GSMs are demonstrated in model organisms *Escherichia coli* and human. Future efforts will be on collecting more single-cell xomics data and curating non-model oragnisms. The XomicsDB can be found at: http://xomicsdb.com.

## Introduction

Life can be defined as a fully autonomous “self-sustaining chemical system capable of Darwinian evolution”, adapted from the version by NASA (https://astrobiology.nasa.gov/research/life-detection/about/) (1). This definition highlights operational spontaneity of life systems in response to internal and external perturbations, governed by the physical and chemical laws. In light of this, the most effective way to study life is through systems biology (2-4), now widely recognized as a new paradigm in biology and a powerful investigative methodology. Systems biology employs computational analyses of complex biological systems, emphasizing holistic operations and interactions among components rather than isolated parts (5-7). A core tenet of this field is the iterative cycle of hypothesis generation, computational modeling, and experimental validation until a comprehensive holistic operation of cellular life is achieved (8).

Central to iterative systems biology is the modeling and integration of multi-omics data based on genome-scale metabolic networks (GSMs) (5,7). Beyond theoretical modeling developments, multi-omics data—referred to here as xomics data—are essential for systems biology research. Moreover, the growing application of AlphaFold3 (9), makes xomics databases even more critical, as paired metabolomics, proteomics, and transcriptomics datasets enable studies of gene regulation in both model and non-model organisms by examining which metabolites bind transcription factors (proteins) and how the metabolite-TF complexes in turn bind DNA to regulate gene expression. However, systems biology imposes strict requirements on xomics datasets. First, different xomics data types must be paired to be useful, meaning each omics data type should be collected from the same batch of biological replicates in the same studies. Second, the organisms for which data are collected should ideally be model organisms, such that extensive structural and functional knowledge is already available for use in systems biology simulations.

Despite the abundance of single-omics datasets, no databases have yet been established specifically to curate and disseminate paired xomics data. One of the closest resources is the Omics Discovery Index (OmicsDI), which aggregates four types of unpaired omics datasets— transcriptomics, proteomics, genomics, and metabolomics—from 11 repositories within a unified structure (9). With the rapid expansion of single-cell multi-omics sequencing, the integration of single-cell xomics data likewise requires high-quality paired datasets.

Here, we present XomicsDB, a resource for curating published paired omics datasets to support systems biology research, with two case studies of xomics data integration. The data are retrieved from published studies at both bulk and single-cell levels. This database will be instrumental for advancing systems biology and related fields, serving as a central hub for curating paired xomics datasets.

### Data Collection

All paired xomics datasets were manually identified and retrieved from published studies, including both bulk and single-cell xomics datasets for model organisms (Supplementary Table 1). Non-model organisms were not considered at the moment but will be included when datasets accumulate for them.

### Database construction

XomicsDB was deployed on an Ubuntu Linux server (version 22), with web services handled by a Nginx reverse proxy. The representation and logic layers were developed using Web 2.0 technologies—HTML5, CSS3, JavaScript (along with the jQuery library)—and PHP for server-side scripting. All data were stored in an optimized MySQL relational database. A keyword-based search engine, powered by the open-source Sphinx Search Server (http://sphinxsearch.com), was integrated into XomicsDB using an HTML inline frame (iframe).

### Datasets statistics

A total of 13 model organisms are represented in the xomics datasets (Figure 1A). Among them, the house mouse (*Mus musculus*) has the largest number of datasets, followed by human and *Escherichia coli* (Figure 1B). In contrast, only a single paired xomics dataset was obtained from the fruit fly (*Drosophila melanogaster*) and the common eastern firefly (*Photinus pyralis*). Notably, most of the datasets were generated from animal species.

**Figure 1.**
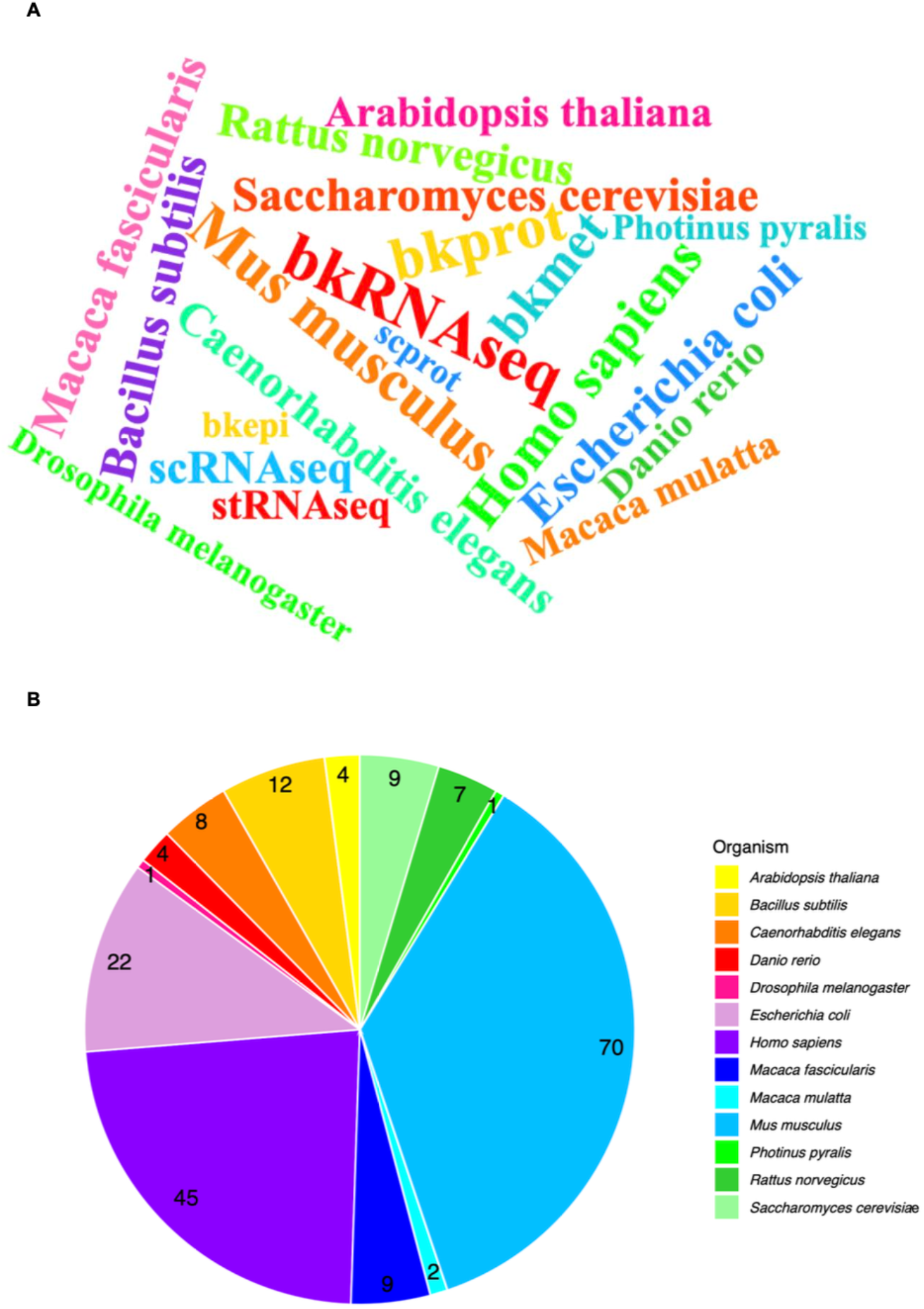
The model organisms of the collected xomics datasets. (A) the word cloud of the organisms for which datasets were collected. (B) the number of datasets for each organism.

As shown in Figure 2, 183 datasets are from bulk xomics experiments, while 17 are from single-cell xomics. Within the bulk category, RNA-seq is the most common (95 datasets), followed by proteomics (59) and metabolomics (28). For single-cell data, RNA-seq accounts for the highest number of datasets (7).

**Figure 2.**
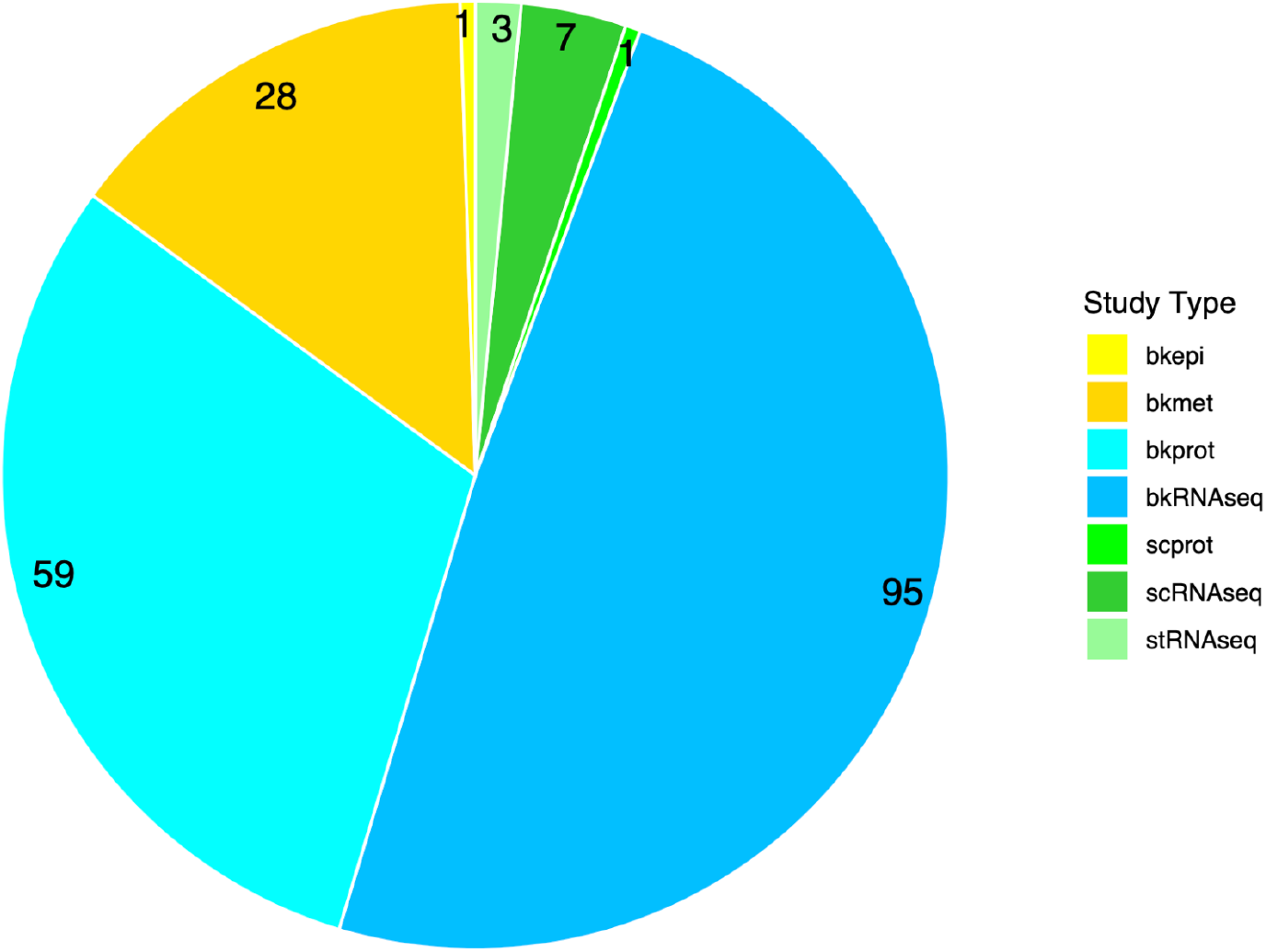
The number of datasets in each omics type. The following abbreviations are used: bkepi represents bulk epigenomics; bkmet, bulk metabolomics; bkprot, bulk proteomics; bkRNAseq, bulk transcriptomics; scprot, single-cell proteomics; scRNAseq, single-cell transcriptomics; and stRNAseq, spatial transcriptomics.

To maximize the utility of xomics datasets, they should ideally be generated from consistent biological replicates within the same study and organism, ensuring greater comparability and reproducibility. So we enumerate the type and number of xomics datasets each organism has. As shown in Figure 3A, the house mouse (*Mus musculus*) which has the most datasets also has the largest number of (six) data types, interestingly followed by human and *Escherichia coli*. The organisms shown to have only one data type are published studies which used paired oxmics data but so far has only made public one of the data types: fruit fly (*D. melanogaster*), the common eastern firefly *P. pyralis, Arabidopsis thaliana*, and the rhesus macaque *Macaca mulatta*. We also provide the specific pairs of xomics data for each of the organisms. Shown here are the two top organisms have the most pairs of xomics data: the house mouse *M. musculus* (Figure 3B) and human (Figure 3C). The rest are shown in supplementary figues (Supplementary Figure 1A-G).

**Figure 3.**
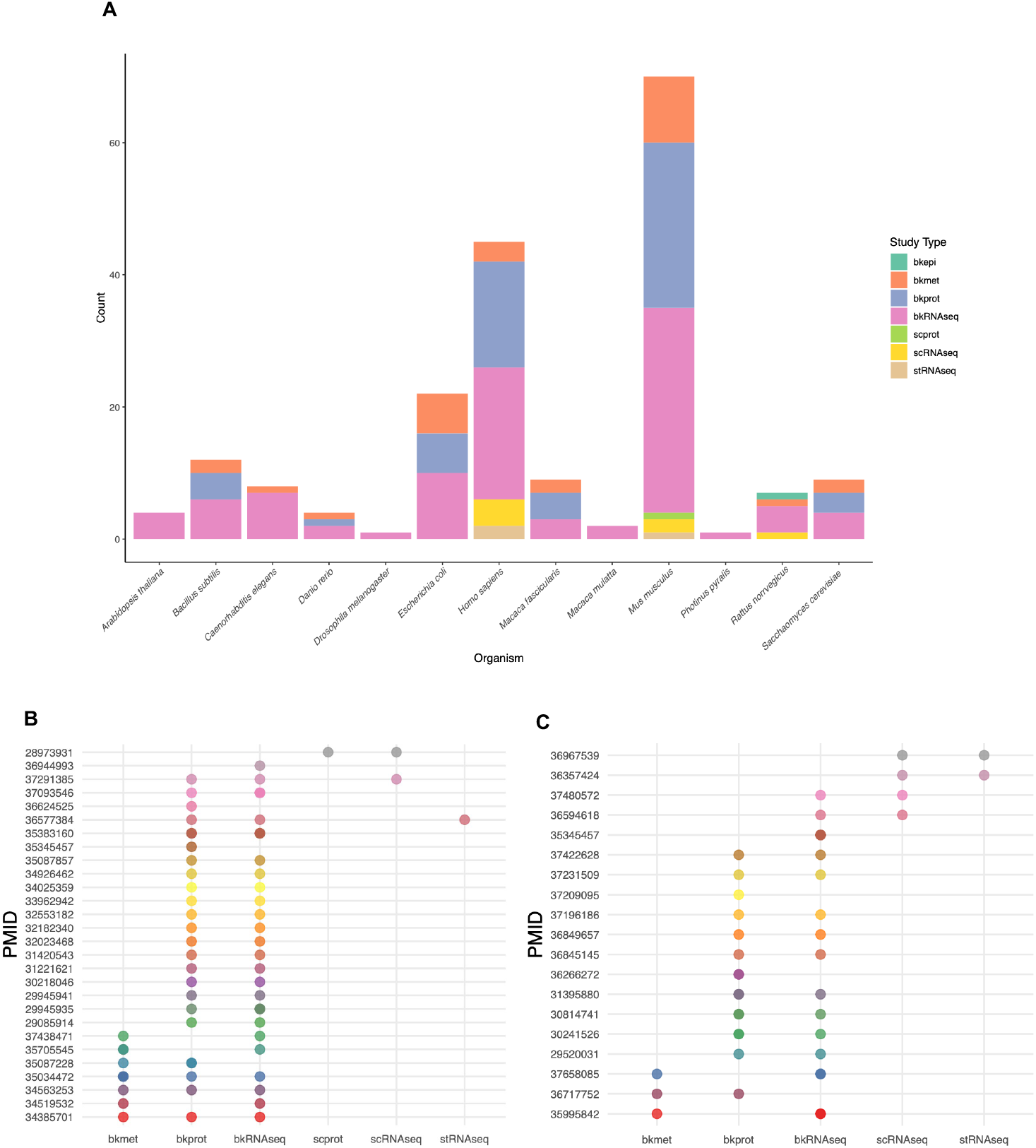
The types of datasets collected in different model organisms. (A) the total of xomics datasets in each organism; (B) the specific pairs of xomics dataset in house mouse (*M. musculus*); and (C) the specific pairs of xomics dataset in human.

### Database Applications

Users can explore the datasets in XomicsDB through four primary entry points: study ID, data type, organism, and accession number/ID. Each browsing method is supported by dedicated pages with integrated search functionality. To promote data sharing, all datasets are available for download, with the exception of a few extremely large proteomics datasets (approximately 1.0 TB each). These larger datasets can be accessed via original source links provided on their respective pages. All other datasets are downloadable either individually from their specific browsing pages or collectively via our FTP server. As we aim to serve as a central repository for paired omics datasets, we encourage users to submit their own paired x-omics data through the dedicated upload page on XomicsDB.

### Case Studies

To demonstrate the utility of paired xomics datasets, here we provide a case study using genome-scale metabolic models (GSMs). GSMs are the natural systems to incorporate xomics data to infer biological operation and regulation. Here we used two model organisms, *Homo sapiens* and *Escherichia coli*, using the GECKO (Genome-scale model to account for Enzyme Constraints using Kinetics and Omics) framework (10). GECKO is selected because it creates an effective way of constraining enzymes by converting them to pseudo-metabolites in GSMs, in which enzyme turnover numbers (*k*_cat_ values) and concentrations (proteomic levels) can be used.

For each organism, two versions of the model were generated: a light model that incorporates only kinetic constraints derived from enzyme turnover rates, and a full model in which these constraints are further refined using proteomics data to limit reaction fluxes based on measured enzyme abundances. The human GSM Human1 was used (11), with protein abundance data from the HCT116 colorectal carcinoma cell line were used to parameterize the full model (12). For *E. coli* grown on glucose minimal medium, the GSM *i*ML1515 for the K-12 MG1655 strain (13) was used and constrained using proteomics data (14).

Flux balance analysis (FBA) in GSMs can compute the steady-state distribution of reaction fluxes and Flux variability analysis (FVA) quantifies the allowable range of fluxes for each reaction under different constraint models, thereby showing how reliable the flux predicitons are. So, FVA was used to showcase the xomics utlitty, by computing the steady-state distribution of reaction fluxes, using biomass production as the objective function for both the *E. coli* and human GSMs.

The comparison of unconstrained, light (constrained with *k*_cat_ values), and full (constrained with *k*_cat_ and proteomics data) models revealed significant differences in metabolic flexibility across the constraint levels. In both the human and *E. coli* models, the application of enzyme constraints through the GECKO framework, particularly the integration of proteomics data in the full model, led to a substantial reduction (more accurate) in the total flux capacity. Flux variability analysis (FVA) showed that the allowable flux ranges became progressively narrower from unconstrained models to full models in both organisms (Fig. 4A–B). In the human model, the median flux range decreased from 17.75 (unconstrained model) to 11.54 (light model) and 0.999 (full model). In *E. coli*, the median dropped from 1000 to 1.34 and 0.091, respectively.

**Figure 4.**
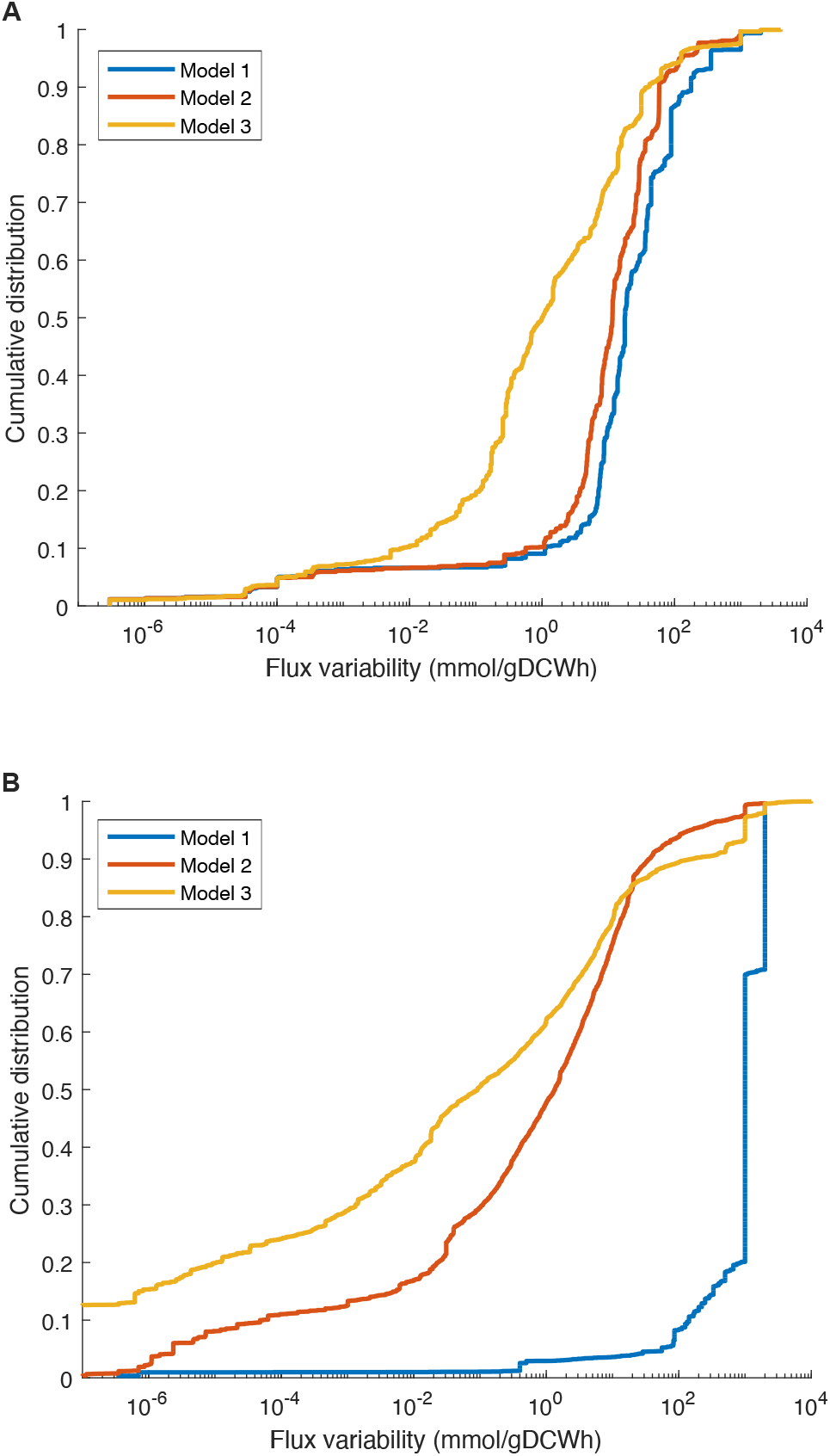
Flux variability analysis for enzyme-constrained models. (A) *Escherichia coli* K-12 metabolic models.(B) Human genome-scale metabolic models. In both panels, Model 1 represents the original unconstrained model; Model 2 is the light enzyme-constrained model incorporating only kinetic constraints (*k*_cat_); and Model 3 is the full enzyme-constrained model incorporating both kinetic and proteomics-based constraints.

## Discussion

Xomics data have been increasingly used in life science studies and now become a common practice. This established paradigm calls for model-based integration of xomics for functional inference, which is the systems biology (2,3,15), the integration of xomics into GSMs (5-7). Such development calls for good-quality paired xomics datasets. The XomicDB developed in this study provides such a data hub for that objective.

This is the first step in curating paired xomics data. Future efforts will continue to collect and curate paired xomics dataset, particularly at the single-cell level. Single-cell xomics data are a bit harder to integrate than bulk xomics due to the existing data types have different throughput in terms of the number of target molecules (e.g., genes, transcripts, cell location, metabolites, proteins, etc.) (16,17). The other bottleneck is slow development in systems biology theories in integrating these single-cell xomics data (18-20).

Xomics from non-model organism will also be collected in the future. As these organisms are not well understood in biology as model organisms, the xomics data may not be efficiently integrated. From a different perspective, these non-model organism are useful in comparison studies with model organisms.

In summary, XomicsDB is a database for paired xomics datasets for model organisms and will be further expanded to include more single-cell xomics and non-model organisms in the future.

## Data Availibility

All data is publicly available for download here: http://xomicsdb.com.

## Supplmentary Data

Supplementary Data are available at NAR online.

## Funding

This work was funded by Kunshan Municipal Government Research Fund [21KNSFC032] ; the National Natural Science Foundation of China [32171565], and DKU Foundation and Chancellor’s Fund [24KDKUF070].

## Conflict of Interest

None.

## Notes

http://xomicsdb.com

### Competing Interest Statement

The authors have declared no competing interest.

