## Supplementary Figure 1 for "XomicsDB: a database of paired multi-omics datasets for systems biology"

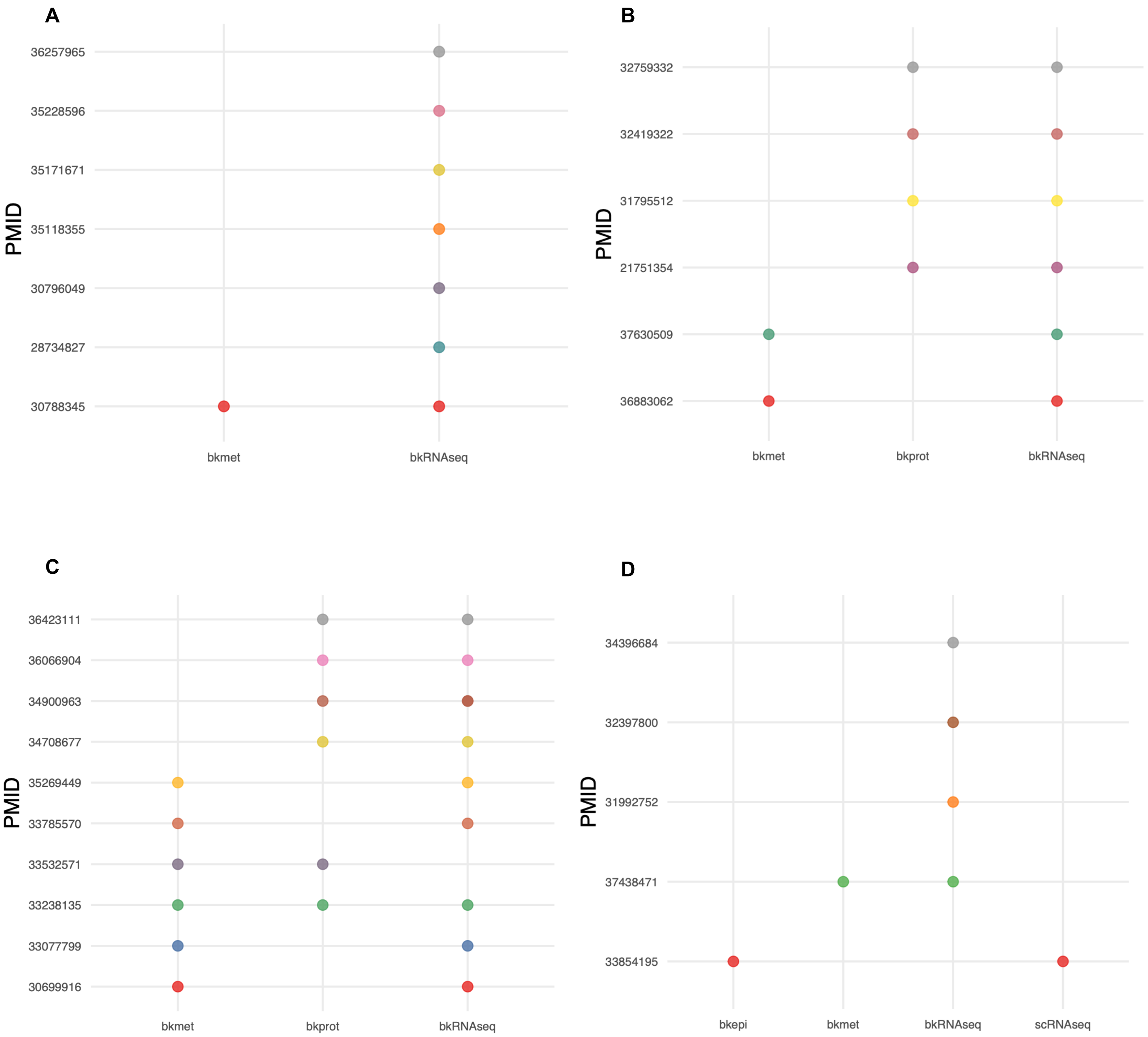
A
PMID
36257965
35228596
35171671
35118355
30796049
28734827
30788345
bkmet
bkRNAseq
B
PMID
32759332
32419322
31795512
21751354
37630509
36883062
bkmet
bkprot
bkRNAseq
C
PMID
36423111
36066904
34900963
34708677
35269449
33785570
33532571
33238135
33077799
30699916
bkmet
bkprot
bkRNAseq
D
PMID
34396684
32397800
31992752
37438471
33854195
bkepi
bkmet
bkRNAseq
scRNAseq

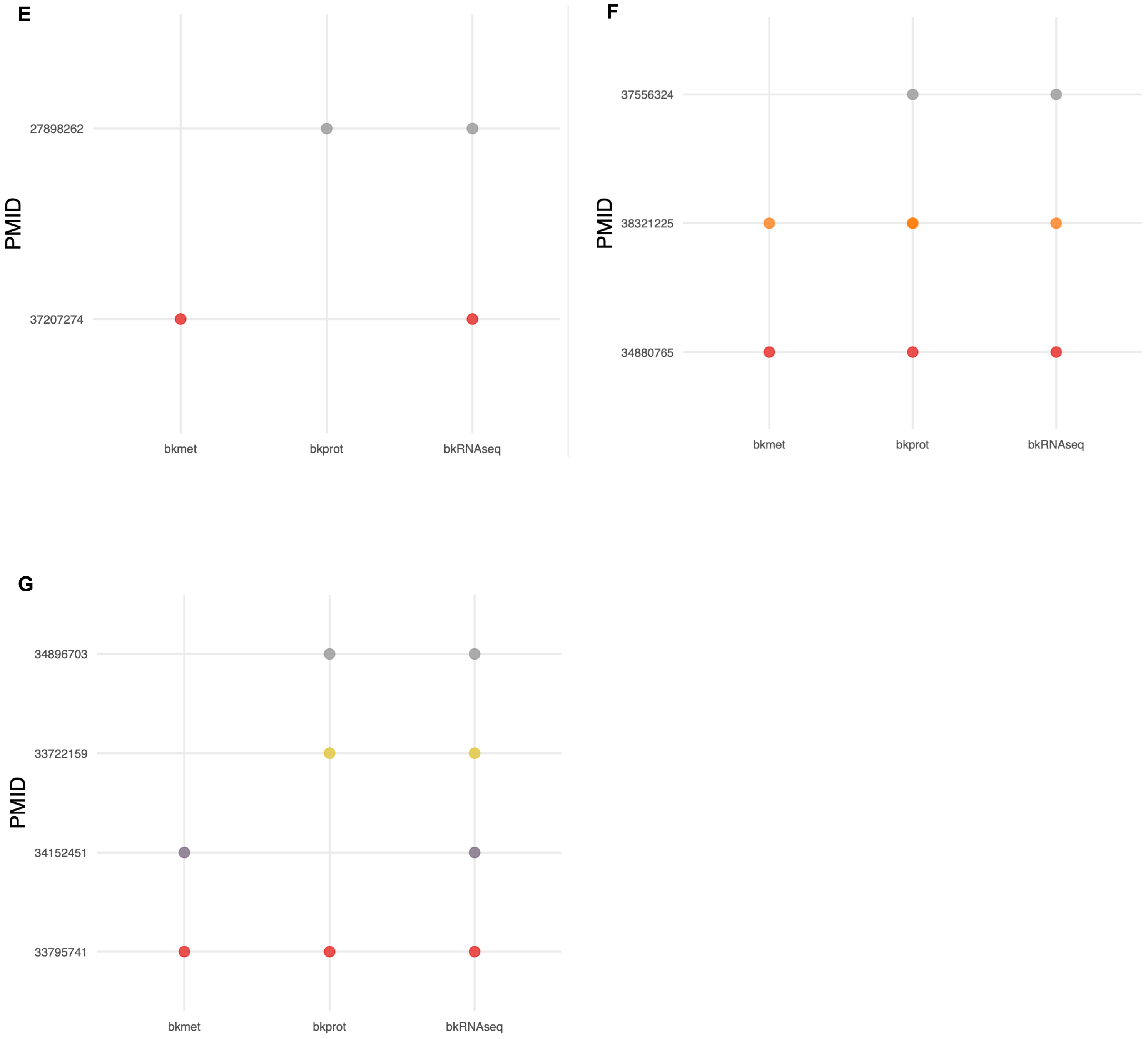


Supplementary Figure 1: Additional paired xomics datasets identified in other model organisms. (A) *Caenorhabditis elegans* ,(B) *Bacillus subtilis.* (C) *Escherichia coli,* (D) *Rattus norvegicus* ,(E) *Danio rerio,*(F) *Macaca fascicularis,*(G) *Saccharomyces cerevisiae*
