## Supplementary Table 1 for "XomicsDB: a database of paired multi-omics datasets for systems biology"

Supplementary table 1: Summary of all included xomics datasets. For each study, the PubMed ID (PMID), study type (bulk or single-cell), and the corresponding model organism are provided.

| **Study ID(PMID)** | **Study Type** | **Organism** |
|---|---|---|
| 21751354 | bulk transcriptomics | *Bacillus subtilis* |
| 21751354 | bulk proteomics | *Bacillus subtilis* |
| 27898262 | bulk proteomics | *Danio rerio* |
| 27898262 | bulk transcriptomics | *Danio rerio* |
| 28004739 | bulk transcriptomics | *Photinus pyralis* |
| 28734827 | bulk transcriptomics | *Caenorhabditis elegans* |
| 28973931 | single-cell transcriptomics | *Mus musculus* |
| 28973931 | single-cell proteomics | *Mus musculus* |
| 29085914 | bulk transcriptomics | *Mus musculus* |
| 29085914 | bulk proteomics | *Mus musculus* |
| 29520031 | bulk transcriptomics | *Homo sapiens* |
| 29520031 | bulk proteomics | *Homo sapiens* |
| 29945935 | bulk transcriptomics | *Mus musculus* |
| 29945935 | bulk transcriptomics | *Mus musculus* |
| 29945935 | bulk transcriptomics | *Mus musculus* |
| 29945935 | bulk transcriptomics | *Mus musculus* |
| 29945935 | bulk proteomics | *Mus musculus* |
| 29945941 | bulk transcriptomics | *Mus musculus* |
| 29945941 | bulk proteomics | *Mus musculus* |
| 30218046 | bulk transcriptomics | *Mus musculus* |
| 30218046 | bulk proteomics | *Mus musculus* |
| 30241526 | bulk transcriptomics | *Homo sapiens* |
| 30241526 | bulk proteomics | *Homo sapiens* |
| 30241526 | bulk proteomics | *Homo sapiens* |
| 30699916 | bulk transcriptomics | *Escherichia coli* |
| 30699916 | bulk metabolomics | *Escherichia coli* |
| 30788345 | bulk transcriptomics | *Caenorhabditis elegans* |
| 30788345 | bulk metabolomics | *Caenorhabditis elegans* |
| 30796049 | bulk transcriptomics | *Caenorhabditis elegans* |

| | | |
|---|---|---|
| 30814741 | bulk transcriptomics | *Homo sapiens* |
| 30814741 | bulk proteomics | *Homo sapiens* |
| 30814741 | bulk proteomics | *Homo sapiens* |
| 31221621 | bulk transcriptomics | *Mus musculus* |
| 31221621 | bulk proteomics | *Mus musculus* |
| 31395880 | bulk transcriptomics | *Homo sapiens* |
| 31395880 | bulk proteomics | *Homo sapiens* |
| 31395880 | bulk proteomics | *Homo sapiens* |
| 31420543 | bulk transcriptomics | *Mus musculus* |
| 31420543 | bulk proteomics | *Mus musculus* |
| 31795512 | bulk transcriptomics | *Bacillus subtilis* |
| 31795512 | bulk proteomics | *Bacillus subtilis* |
| 31992752 | bulk transcriptomics | *Rattus norvegicus* |
| 32023468 | bulk transcriptomics | *Mus musculus* |
| 32023468 | bulk proteomics | *Mus musculus* |
| 32170290 | bulk transcriptomics | *Arabidopsis thaliana* |
| 32182340 | bulk transcriptomics | *Mus musculus* |
| 32182340 | bulk proteomics | *Mus musculus* |
| 32397800 | bulk transcriptomics | *Rattus norvegicus* |
| 32419322 | bulk transcriptomics | *Bacillus subtilis* |
| 32419322 | bulk proteomics | *Bacillus subtilis* |
| 32483184 | bulk transcriptomics | *Drosophila melanogaster* |
| 32553182 | bulk transcriptomics | *Mus musculus* |
| 32553182 | bulk proteomics | *Mus musculus* |
| 32759332 | bulk transcriptomics | *Bacillus subtilis* |
| 32759332 | bulk proteomics | *Bacillus subtilis* |
| 33077799 | bulk transcriptomics | *Escherichia coli* |
| 33077799 | bulk metabolomics | *Escherichia coli* |
| 33238135 | bulk proteomics | *Escherichia coli* |
| 33238135 | bulk metabolomics | *Escherichia coli* |
| 33238135 | bulk transcriptomics | *Escherichia coli* |
| 33532571 | bulk proteomics | *Escherichia coli* |

| | | |
|---|---|---|
| 33532571 | bulk metabolomics | *Escherichia coli* |
| 33722159 | bulk transcriptomics | *Saccharomyces cerevisiae* |
| 33722159 | bulk proteomics | *Saccharomyces cerevisiae* |
| 33785570 | bulk transcriptomics | *Escherichia coli* |
| 33785570 | bulk metabolomics | *Escherichia coli* |
| 33795741 | bulk metabolomics | *Saccharomyces cerevisiae* |
| 33795741 | bulk proteomics | *Saccharomyces cerevisiae* |
| 33795741 | bulk transcriptomics | *Saccharomyces cerevisiae* |
| 33854195 | single-cell transcriptomics | *Rattus norvegicus* |
| 33854195 | bulk epigenomics | *Rattus norvegicus* |
| 33962942 | bulk transcriptomics | *Mus musculus* |
| 33962942 | bulk proteomics | *Mus musculus* |
| 34025359 | bulk proteomics | *Mus musculus* |
| 34025359 | bulk transcriptomics | *Mus musculus* |
| 34152451 | bulk metabolomics | *Saccharomyces cerevisiae* |
| 34152451 | bulk transcriptomics | *Saccharomyces cerevisiae* |
| 34385701 | bulk proteomics | *Mus musculus* |
| 34385701 | bulk metabolomics | *Mus musculus* |
| 34385701 | bulk transcriptomics | *Mus musculus* |
| 34396684 | bulk transcriptomics | *Rattus norvegicus* |
| 34519532 | bulk metabolomics | *Mus musculus* |
| 34519532 | bulk transcriptomics | *Mus musculus* |
| 34563253 | bulk transcriptomics | *Mus musculus* |
| 34563253 | bulk proteomics | *Mus musculus* |
| 34563253 | bulk metabolomics | *Mus musculus* |
| 34708677 | bulk proteomics | *Escherichia coli* |
| 34708677 | bulk transcriptomics | *Escherichia coli* |
| 34880765 | bulk transcriptomics | *Macaca fascicularis* |
| 34880765 | bulk proteomics | *Macaca fascicularis* |

| | | |
|---|---|---|
| 34880765 | bulk metabolomics | *Macaca fascicularis* |
| 34896703 | bulk proteomics | *Saccharomyces cerevisiae* |
| 34896703 | bulk transcriptomics | *Saccharomyces cerevisiae* |
| 34900963 | bulk transcriptomics | *Escherichia coli* |
| 34900963 | bulk proteomics | *Escherichia coli* |
| 34900963 | bulk transcriptomics | *Escherichia coli* |
| 34926462 | bulk proteomics | *Mus musculus* |
| 34926462 | bulk transcriptomics | *Mus musculus* |
| 35034472 | bulk transcriptomics | *Mus musculus* |
| 35034472 | bulk proteomics | *Mus musculus* |
| 35034472 | bulk metabolomics | *Mus musculus* |
| 35034472 | bulk metabolomics | *Mus musculus* |
| 35034472 | bulk metabolomics | *Mus musculus* |
| 35087228 | bulk proteomics | *Mus musculus* |
| 35087228 | bulk proteomics | *Mus musculus* |
| 35087228 | bulk metabolomics | *Mus musculus* |
| 35087857 | bulk proteomics | *Mus musculus* |
| 35087857 | bulk transcriptomics | *Mus musculus* |
| 35118355 | bulk transcriptomics | *Caenorhabditis elegans* |
| 35171671 | bulk transcriptomics | *Caenorhabditis elegans* |
| 35228596 | bulk transcriptomics | *Caenorhabditis elegans* |
| 35269449 | bulk transcriptomics | *Escherichia coli* |
| 35269449 | bulk metabolomics | *Escherichia coli* |
| 35345457 | bulk transcriptomics | *Homo sapiens* |
| 35345457 | bulk transcriptomics | *Homo sapiens* |
| 35345457 | bulk transcriptomics | *Homo sapiens* |
| 35345457 | bulk transcriptomics | *Homo sapiens* |
| 35345457 | bulk transcriptomics | *Homo sapiens* |
| 35345457 | bulk proteomics | *Mus musculus (Human cell injected into the mouse)* |
| 35383160 | bulk transcriptomics | *Mus musculus* |

| | | |
|---|---|---|
| 35383160 | bulk transcriptomics | *Mus musculus* |
| 35383160 | bulk transcriptomics | *Mus musculus* |
| 35383160 | bulk proteomics | *Mus musculus* |
| 35383160 | bulk proteomics | *Mus musculus* |
| 35435236 | bulk transcriptomics | *Arabidopsis thaliana* |
| 35435236 | bulk transcriptomics | *Arabidopsis thaliana* |
| 35705545 | bulk transcriptomics | *Mus musculus* |
| 35705545 | bulk metabolomics | *Mus musculus* |
| 35705545 | bulk metabolomics | *Mus musculus* |
| 35892179 | bulk transcriptomics | *Arabidopsis thaliana* |
| 35995842 | bulk transcriptomics | *Homo sapiens* |
| 35995842 | bulk transcriptomics | *Homo sapiens* |
| 35995842 | bulk metabolomics | *Homo sapiens* |
| 36066904 | bulk transcriptomics | *Escherichia coli* |
| 36066904 | bulk proteomics | *Escherichia coli* |
| 36130949 | bulk transcriptomics | *Macaca mulatta* |
| 36257965 | bulk transcriptomics | *Caenorhabditis elegans* |
| 36266272 | bulk proteomics | *Homo sapiens* |
| 36266272 | bulk proteomics | *Homo sapiens* |
| 36357424 | single-cell transcriptomics | *Homo sapiens* |
| 36357424 | spatial transcriptomics | *Homo sapiens* |
| 36423111 | bulk proteomics | *Escherichia coli* |
| 36423111 | bulk transcriptomics | *Escherichia coli* |
| 36577384 | bulk transcriptomics | *Mus musculus* |
| 36577384 | spatial transcriptomics | *Mus musculus* |
| 36577384 | bulk proteomics | *Mus musculus* |
| 36594618 | bulk transcriptomics | *Homo sapiens* |
| 36594618 | single-cell transcriptomics | *Homo sapiens* |
| 36624525 | bulk proteomics | *Mus musculus* |
| 36717752 | bulk proteomics | *Homo sapiens* |
| 36717752 | bulk metabolomics | *Homo sapiens* |
| 36845145 | bulk transcriptomics | *Homo sapiens* |

| | | |
|---|---|---|
| 36845145 | bulk proteomics | *Homo sapiens* |
| 36849657 | bulk transcriptomics | *Homo sapiens* |
| 36849657 | bulk proteomics | *Homo sapiens* |
| 36883062 | bulk metabolomics | *Bacillus subtilis* |
| 36883062 | bulk transcriptomics | *Bacillus subtilis* |
| 36944971 | bulk transcriptomics | *Macaca mulatta* |
| 36944993 | bulk transcriptomics | *Mus musculus* |
| 36967539 | single-cell transcriptomics | *Homo sapiens* |
| 36967539 | spatial transcriptomics | *Homo sapiens* |
| 37093546 | bulk transcriptomics | *Mus musculus* |
| 37093546 | bulk transcriptomics | *Mus musculus* |
| 37093546 | bulk transcriptomics | *Mus musculus* |
| 37093546 | bulk proteomics | *Mus musculus* |
| 37196186 | bulk transcriptomics | *Homo sapiens* |
| 37196186 | bulk proteomics | *Homo sapiens* |
| 37207274 | bulk metabolomics | *Danio rerio* |
| 37207274 | bulk transcriptomics | *Danio rerio* |
| 37209095 | bulk proteomics | *Homo sapiens* |
| 37231509 | bulk transcriptomics | *Homo sapiens* |
| 37231509 | bulk proteomics | *Homo sapiens* |
| 37291385 | bulk transcriptomics & single-cell transcriptomics | *Mus musculus* |
| 37291385 | bulk proteomics | *Mus musculus* |
| 37422628 | bulk transcriptomics | *Homo sapiens* |
| 37422628 | bulk proteomics | *Homo sapiens* |
| 37438471 | bulk transcriptomics | *Rattus norvegicus,Mus musculus* |
| 37438471 | bulk metabolomics | *Rattus norvegicus, Mus musculus* |
| 37480572 | bulk transcriptomics | *Homo sapiens* |
| 37480572 | single-cell transcriptomics | *Homo sapiens* |
| 37556324 | bulk proteomics | *Macaca fascicularis* |
| 37556324 | bulk transcriptomics | *Macaca fascicularis* |

| | | |
|---|---|---|
| 37630509 | bulk transcriptomics | *Bacillus subtilis* |
| 37630509 | bulk metabolomics | *Bacillus subtilis* |
| 37658085 | bulk transcriptomics | *Homo sapiens* |
| 37658085 | bulk transcriptomics | *Homo sapiens* |
| 37658085 | bulk metabolomics | *Homo sapiens* |
| 38321225 | bulk proteomics | *Macaca fascicularis* |
| 38321225 | bulk metabolomics | *Macaca fascicularis* |
| 38321225 | bulk transcriptomics | *Macaca fascicularis* |
| 38321225 | bulk proteomics | *Macaca fascicularis* |
